# Meiotic recombination in *Leptosphaeria maculans* favours the rise of transgressive isolates adapted to a nonhost species, *Brassica carinata*

**DOI:** 10.1101/2024.02.15.580423

**Authors:** Julie M. Noah, Mathilde Gorse, Carole-Anne Romain, Elise J. Gay, Thierry Rouxel, Marie-Hélène Balesdent, Jessica L. Soyer

## Abstract

*Leptosphaeria maculans* is one of the major fungal pathogens on oilseed rape (*Brassica napus*), causing stem canker disease. The closely related *Brassica* species *Brassica nigra*, *Brassica juncea,* and *Brassica carinata* display extreme resistance toward stem canker. In this study, we demonstrate the nonhost status of *B. carinata* toward *L. maculans* in France through field experiments and inoculations performed in controlled conditions. A few isolates moderately adapted to *B. carinata*, in controlled conditions, were recovered in the field on *B. nigra* leaves, allowing us to investigate the unusual *B. carinata* – *L. maculans* interactions using molecular, macroscopic, and microscopic analyses. A cross between a *L. maculans* isolate adapted to *B. napus* and an isolate moderately adapted to *B. carinata* allowed the generation, in the lab, of recombinant *L. maculans* strains better adapted to *B. carinata* than the natural parental isolate obtained from *B. nigra*, and highlighted the polygenic determinism of the adaptation of *L. maculans* to *B. carinata* and *B. napus*. This biological material will allow further investigation of the molecular determinants of the adaptation of *L. maculans* to nonhost species and elucidate the genetic resistance basis of *B. carinata*.

## Introduction

Phytopathogenic fungi are one of the major problems in agriculture, causing economic and food production problems (Pennesi, 2010). However, most interactions between a potential plant pathogen and a potential host plant do not lead to disease. Plants exhibit two types of disease resistance: nonhost resistance (NHR) (Heath, 2000) and host-specific resistance, with NHR being the most widespread as it prevents most plants from being infected by potential pathogens. In NHR, all genotypes of a considered plant species are resistant to all genotypes of a potential pathogen species. Host resistance is genotype-specific, as some genotypes of a given plant species are resistant toward some genotypes of the adapted pathogen species (Niks and Marcel, 2009). Mechanisms responsible for NHR remain to be fully deciphered, but some resistance mechanisms involved in host-specific and nonhost resistances converge (Gill *et al*., 2015). In NHR, two types of resistance can be distinguished: type-I and type-II nonhost resistances (Schulze-Lefert and Panstruga, 2011). Type-I NHR is a pre-invasive resistance mechanism characterized by a complete absence of visible symptoms, thus involving passive defence plant mechanisms. On the contrary, type-II NHR, a post-invasive resistance mechanism, is characterized by a local cell death, called hypersensitive response (HR; Gill *et al*., 2015). While the defence mechanisms set up in host-specific or nonhost resistance can involve the same actors, the difference between both mechanisms lies in the level at which the resistance is expressed (eitherall genotypes of a given species are resistant or only some genotypes are). To date, NHR is considered more durable than host-specific resistance in the management of plant diseases (Lee *et al*., 2017; Panstruga and Moscou, 2020) and many examples can be found of interspecies transfer of NHR genes into host-species in breeding programs (Zhao *et al*., 2005; Kawashima *et al*., 2016; Wang *et al*., 2018; see more examples in the review of Ayliffe and Sørensen, 2019).

*Leptosphaeria maculans* is a Dothideomycete fungus that causes stem canker of oilseed rape (*Brassica napus*). It is one of the main fungal pathogens of *B. napus* leading to important yield losses, c.a. 20% worldwide (Fitt *et al*., 2006; Rouxel *et al*., 2011). This hemibiotrophic pathogen has a complex life cycle tightly associated with that of its host plant, during which it alternates between different nutrition modes (Rouxel and Balesdent, 2005). Firstly, it survives and develops as a saprophyte on infected crop residues, on which sexual reproduction occurs. Ascospores are then disseminated by the wind and infect seedlings (e.g. in autumn on winter oilseed rape in Europe) after the penetration of hyphae through stomata or natural wounds (Hammond and Lewis, 1987). The asymptomatic development in young leaves is followed by a short necrotrophic phase, resulting in the development of greyish spots on leaves leading to asexual reproduction. *L. maculans* then initiates systemic and biotrophic colonisation from the petiole to the stem basis, progressing in and around the vascular bundles (Hammond *et al*., 1985). After several months of asymptomatic colonization in the stem, necrosis appears at the stem basis, causing plant lodging and resulting in yield losses (Rouxel and Balesdent, 2005). In the field, *L. maculans* is often associated with the closely related species *Leptosphaeria biglobosa* (West *et al*., 2001), and these two Leptosphaeria species can be found together on the same host and even in the same plant tissues (Jacques *et al*., 2021). They both cause necrotic lesions and colonize host plant tissues, but their impacts on crops are different as it is postulated that only *L. maculans* can cause lodging and damage to the crops (West *et al*., 2001).

The favoured control method against stem canker of oilseed rape relies on the identification of effective host resistance sources (mostly resistance genes) and the breeding of resistant cultivars. This genetic resistance mechanism relies on a monogenic determinism following the gene-for-gene relationship (Flor, 1955): interaction of the product of a host-specific resistance gene (R) with the product of an avirulence gene (AVR) causes HR, which thus blocks the progression of the pathogen. Until now, 21 specific major resistance genes (*Rlm*), obeying the gene-for-gene relationship, have been identified in different *Brassica* species, and some have been introduced into *B. napus* (Chèvre *et al*., 1997; Ansan-Melayah *et al*., 1998; Delourme *et al*., 2004, 2006; Yu *et al*., 2005, 2008; Rimmer, 2006; Long *et al*., 2011; Degrave *et al*., 2021; Raman *et al*., 2021; see Cantila *et al*., 2021 for a detailed list of the identified resistance genes toward blackleg *Brassica* spp). For instance, *B. rapa* and *B. juncea* have been used in breeding programs in various countries (Raman *et al*., 2013; Cantila *et al*., 2021) to select resistance genes and introduce them into oilseed rape. Nevertheless, *L. maculans* has a great capacity to adapt, for example via the loss, inactivation, or point mutations of avirulence genes leading to an absence of recognition of the avirulence protein by the host surveillance machinery (Rouxel and Balesdent, 2017). This enables *L. maculans* populations to overcome resistance deployed in the field (Daverdin *et al*., 2012; Zhang *et al*., 2015), illustrating the need to find new sources of resistance that would be more durable than resistances currently used in the field. *L. maculans* is a specialist fungus with a limited host range. This fungus can infect a few Brassicaceae species including species from the U triangle (Nagaharu, 1935). The U triangle is a schematic representation of the interspecific hybridization between three ancestral *Brassica* species: *Brassica rapa* (turnip, Chinese cabbages; A genome), *Brassica nigra* (black mustard; B genome), and *Brassica oleracea* (cabbages; C genome). These three diploid species are natural progenitors of the three allotetraploid *Brassica* species: *Brassica napus* (rapeseed, canola; AC genome), *Brassica juncea* (Indian mustard; AB genome), and *Brassica carinata* (Ethiopian mustard; BC genome) (Nagaharu, 1935). The species *B. napus*, *B. oleracea*, and *B. rapa* are susceptible host species of *L. maculans* (Gugel *et al*., 1990). On the contrary, much rarer examples are available in the literature of isolates able to attack *B. juncea*, *B. nigra* or *B. carinata* (Roy, 1978; Gugel *et al*., 1990; Zhu *et al*., 1993; Johnson and Lewis, 1994; Somda *et al*., 1999; Li *et al*., 2005). In analyses from Gugel *et al*. (1990), Johnson and Lewis (1994), Somda *et al*. (1995) or Li *et al*. (2005), isolates able to infect *B. carinata* corresponded to “weakly” virulent isolates of *L. maculans*; these “weakly” virulent isolates were later identified as isolates from the *L. biglobosa* species (Shoemaker and Brun, 2001). However, two *L. maculans* Australian isolates, isolated from *B. napus* in the field, were reported moderately virulent on one *B. carinata* genotype (Li *et al*., 2005).

We analyse here the extreme resistance of *B. carinata* toward *L. maculans* by i) evaluating the field behaviour of *B. carinata* in French conditions and ii) evaluating the adaptive ability of *L. maculans* to this species using phytopathological, cytological, genetic, molecular, and transcriptomic approaches. A collection of natural *L. maculans* isolates, including isolates collected on *B. nigra* (Somda and Brun, 1995) were then evaluated on several *B. napus* and *B. carinata* genotypes to assess their capacity to infect these species. One of the isolates, HB10.19, causing limited symptoms on *B. carinata* cotyledons, was used to screen a large collection of accessins of *B. carinata*. A cross between the reference isolate of *L. maculans*, JN3, causing virulence symptoms on *B. napus* while non-adapted to *B. carinata*, and HB10.19 was done. The progeny isolates displayed a continuum of aggressiveness on cotyledons of both *B. napus* and *B. carinata*, suggesting a complex and multi-genic determinism of the adaptation to the hosts. A few progeny isolates had a transgressive behaviour and could infect both *B. napus* and *B. carinata* cotyledons in controlled conditions. However, these could not proceed to the stems in which they only caused minor symptoms. Altogether, these data indicate that *B. carinata* is a nonhost species of *L. maculans*; they nevertheless highlight that recombination of traits at meiosis can allow the emergence of transgressive *L. maculans* isolates able to infect both *B. napus* and *B. carinata* cotyledons.

## Experimental procedures

### Plant and fungal material

In preliminary experiments, one *B. carinata* line, SD199947, provided by Dr. Silva-Dias, was identified as resistant to a selection of *L. maculans* isolates. Sixteen plants of this line were further self-pollinated in bulk to produce the *B. carinata* line D5.6.12. A collection of 52 accessions of *B. carinata* genotypes (Table S1) were retrieved from the IPK collection (IPK, Gatersleben, Germany) and evaluated at the seedling stage for their resistance toward *L. maculans* isolates. A total of 53 *L. maculans* isolates, present in the INRAE collection of isolates, and from diverse plant and geographical origins was used in pathogenicity tests (Table S2).

### Field experiments

Experimental fields were set up for five cropping seasons as described in Degrave *et al*. (2021) and Jacques *et al*. (2021). In 2008-2009, 2009-2010, 2010-2011, the *B. carinata* line D5.6.12*, B. napus* cv. Goéland (susceptible to French *L. maculans* populations), and the resistant *B. napus* cv. Fu-Eurol-*Rlm10* (in which the *B. nigra* resistance gene *Rlm10* was introgressed) were sown at eight locations in France (Degrave *et al*., 2021). In 2016-2017 and 2017-2018, the genotypes were sown at the INRAE experimental field station at Grignon, France (Jacques *et al*., 2021). Fields were regularly surveyed for the development of *L. maculans* or *L. biglobosa* symptoms and leaf spots were collected to recover *Leptosphaeria* isolates.

### Isolates (culture, sporulation, *in vitro* cross)

*Leptosphaeria* isolates were recovered from single pycnidia on symptoms developed on *Brassica* leaf collected in field experiments as described in Balesdent *et al*. (2023). Isolates of *L. maculans* and *L. biglobosa* were grown on V8-agar at 24°C for sporulation, and stored for long term at 4°C on 1% malt-2% agar slant tubes to establish a collection of isolates. Mycelium of the isolates was collected and placed in 2 ml Eppendorf® tubes for further DNA extraction. An *in vitro* cross between the isolates JN3 and HB10.19 was performed as previously described (Balesdent *et al*., 2001). Progeny of the cross was recovered as described in Plissonneau *et al*. (2016). Fungal cultures and conidia production were performed as described in Ansan-Melayah *et al*. (1995)

### Pathogenicity assays

Pathogenicity assays at the cotyledon stage were performed as described in Balesdent *et al*. (2001), on 12-days-old seedlings of genotypes of *B. napus* and genotypes of *B. carinata*. Two methods were used for inoculation, with or without wounding of the cotyledons. In the former, cotyledons were wounded by a needle and 10 µl of a 10^7^/ml pycnidiospores suspension was put on the wound; in the latter, Tween 20 was added in the pycnidiospore suspension (0.1% [v/v] Tween® 20; Roth, Karlsruhe, Germany) and the suspension was applied with a 0.6-mm brush on the upper part of the cotyledon. After 48 hours of incubation under high humidity and darkness, plants were transferred in a climate chamber at 24°C (day) and 16°C (night) with a photoperiod of 16 hours of light and 80% relative air humidity. Symptoms were scored following the IMASCORE rating scale (Balesdent *et al*., 2001) nine and 12-13 days post-inoculation (dpi); the symptom scale ranges from 1 to 6 with 1-3 in which the fungus is avirulent (hence the plant is resistant), and 4-6 in which the fungus is virulent (hence the plant is susceptible).

Cotyledon inoculation with ascospores from naturally infected stubbles was done using the ascospore shower method. Naturally infected stubbles from susceptible *B. napus* cultivars bearing mature pseudothecia were humidified and placed over 10-days old plants of *B. napus* or *B. carinata* for 48h in the dark. Then, the plants were transferred to growth chambers as described above. Symptoms were surveyed during two weeks in order to recover isolates from symptoms, when present (Plissonneau *et al*., 2017).

For the analysis of stem infection, pathogenicity assays were performed as described in Gervais *et al*. (2017). Three isolates (JN3, HB10.19, and V77.1.11) were used to inoculate the stem of *B. napus* (cv. Yudal) and *B. carinata* (line D5.6.12). For each isolate, 16 plants of each species were inoculated. Plants were inoculated at the three-leaf stage (in our experiment,2 days post-sowing), the petioles were horizontally cut 1 cm from the leaf insertion and 10 µl of a 10^7^/ml pycnidiospores suspension was put on the wound. After 48 hours of incubation under high humidity and darkness, plants were transferred in a climate chamber at 24°C (day) and 16°C (night) with a photoperiod of 16 hours and 80% relative air humidity. The length of the external necrosis of the stem were measured 38 dpi with a calliper then the stem basis was longitudinally cut 5 cm long with a scalpel checking for sign of inner necrosis.

### Identification of the species from leaf symptoms using amplification and sequencing of the ITS locus

20 mg of lyophilised mycelium was used for DNA extraction as described in Jacques *et al*. (2021). Characterization of isolates was based on PCR amplification and sequencing of the Internal transcribed spacer (ITS) locus (Mendes-Pereira *et al*., 2003; Table S3), allowing their assignation to *L. maculans* or *L. biglobosa* species.

### Biomass quantification analysis

Fungal biomass in infected cotyledons of *B. carinata* (line D5.6.12) and *B. napus* (cv. Yudal) was estimated as described in Jacques *et al*. (2021). Five, seven, nine, and 12 dpi, a 0.25-cm² area of around 50 infected plant material was collected around the inoculation point and pooled in a falcon tube, flash frozen in liquid nitrogen, and stored at -80°C until further analysis. The infected material was ground before DNA extraction and 80 to 100 mg of material was used for DNA extraction as described by Jacques *et al*. (2021). For each condition tested, two DNA extractions of two independent biological samples were done. *L. maculans* species-specific qPCR were performed using primers to amplify *LmEF1-α* and a probe (Table S3) as described in Jacques *et al*. (2021). Calibration ranges using DNA from the JN3 isolate as a reference DNA were performed from 5 ng to 5.10^-3^ ng in triplicate, as described in Jacques *et al*. (2021).

### Cytological assays

Cotyledons from two plants per interactions were sampled at seven and 15 dpi following wounded inoculation and maintained in glass vials. Three ml of trypan blue lactophenol solution prepared as follow (0.05% [w/v], 0.02 g trypan blue, 10 ml lactic acid (85% w:w), 10 ml phenol (TE buffer equilibrated, pH 7.5-8.0), 10 ml glycerol (>=99%), 10 ml distilled water) was added to the sample and placed in boiling water for three minutes. The trypan blue solution was then discarded and replaced with a chloral hydrate solution (3 ml of a 2.5 g/ml solution); this solution was changed every three days with fresh chloral hydrate solution until the solution and tissue became colourless. The samples were mounted between slides in 50% glycerol before observation with the Leica DM5550 microscope.

### RNA-seq data and analysis

Naturally infected leaves of *B. napus* (cv. Darmor) and *B. carinata* (line D5.6.12) were collected on November 16 and 29, 2017, from the experimental field at Grignon, France. Symptoms were classified as “atypical”, i.e. presumably due to *L. biglobosa*, “typical”, i.e. presumably due to *L. maculans*, or asymptomatic. Leaf discs (one cm diameter) were collected from the samples, centred on symptoms (or randomly from asymptomatic leaf tissues), for RNA-seq analysis, as described by Gay *et al*. (2021).

Raw RNA-seq reads were mapped with the STAR software version 020201 (Dobin *et al*., 2013) on the reference genomes of *L. maculans* (isolate JN3) and *L. biglobosa* (isolate G12-14) (Dutreux *et al*., 2018). As isolates of *L. biglobosa* and *L. maculans* can be present simultaneously in the same samples, the mapping parameters have been optimized as follows: we chose a maximum number of mismatches of two and a mapping on a concatenated genome of both species. According to the maximum intron size found in the genomes, we allowed an intron size of 10,000 pb. Other parameters have been used as follows: outFilterMultimapNmax: 100; SeedSearchStartLmax: 12; alignSJoverhangMin: 15; alignIntronMin: 10. Then, we selected the properly paired reads in BAM files with Samtools v1.6 (Li *et al*., 2009; Gay *et al*., 2023).

## Results

### *Brassica carinata* displays an extreme resistance to *Leptosphaeria maculans*

Attempts were performed to recover *L. maculans* isolates from *B. carinata* leaves in the field for three consecutive years at eight sites in France. While *L. maculans* was easily isolated from leaf spots on both susceptible or resistant *B. napus* genotypes –respectively 98% and 85.5% of the isolates-, mainly *L. biglobosa* (97.5%) isolates were recovered from *B. carinata* leaf symptoms. Furthermore, the two *L. maculans* isolates recovered from *B. carinata* were not able to produce symptoms on *B. carinata* after re-inoculation in controlled conditions, while 56.6% of the isolates recovered from *B. napus* with the specific resistance gene *Rlm10* were found virulent after inoculation on *B. napus-Rlm10* (Table 1). Isolates were also recovered from symptoms developed on *B. carinata* leaves during two other consecutive years (autumn 2016 and 2017) in fields at Grignon, France (Table 1; Fig. S1). Among the isolates recovered on *B. carinata*, one corresponded to *L. maculans*, and all the others to *L. biglobosa*. The *L. maculans* isolate was unable to induce symptoms on *B. carinata* cotyledons under controlled conditions (Table 1).

**Table 1.**
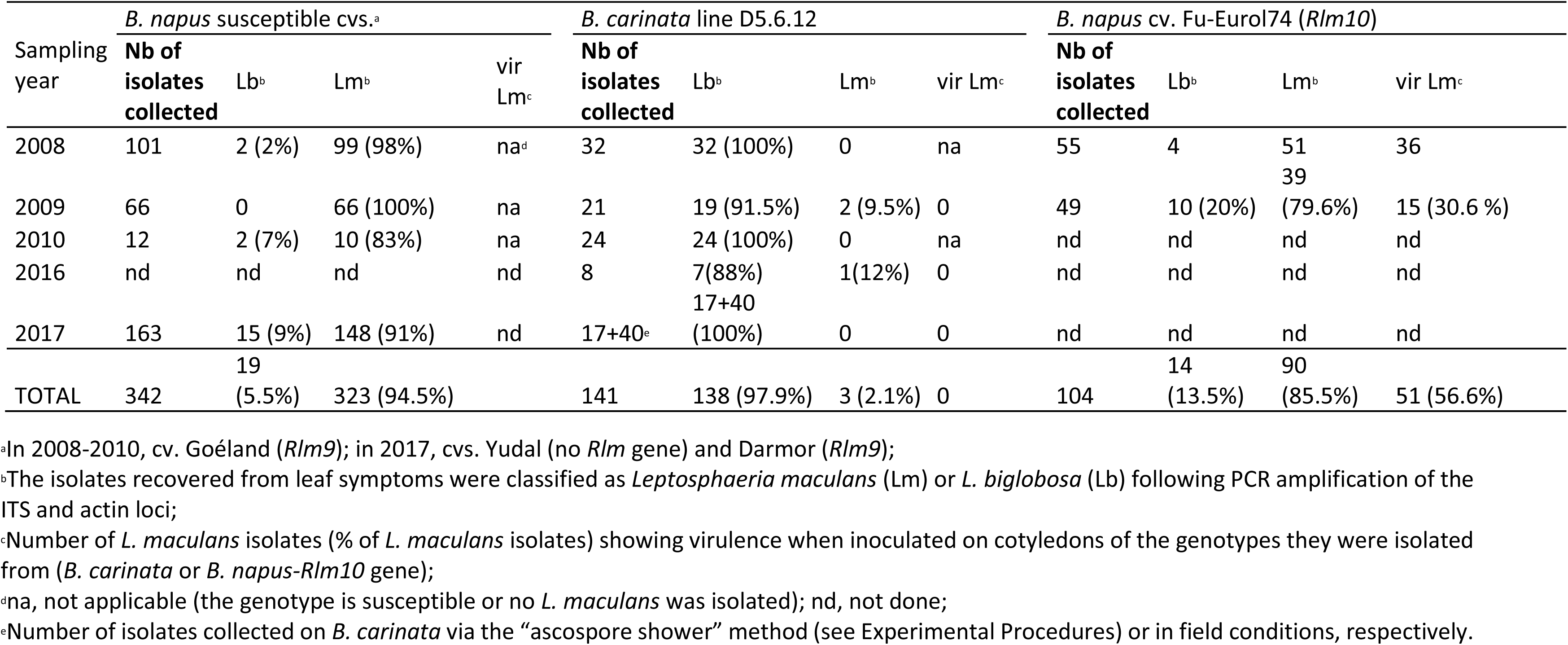
Comparison of *Leptosphaeria* populations recovered from *Brassica napus* and *Brassica carinata* under field conditions.

An additional attempt was performed to recover *L. maculans* isolates on *B. carinata* with the “ascospore shower method”. This method allowed us to place plantlets of *B. carinata* under residues of oilseed rape bearing mature pseudothecia. Ascospores released from the pseudothecia produced leaf spots when ejected on *B. napus* plants, but no typical *L. maculans* symptoms appeared on the *B. carinata* D5.6.12 line. Seventeen isolates were nevertheless recovered from the symptoms developed on cotyledons of *B. carinata*. All these isolates corresponded to *L. biglobosa* (Table 1) and induced avirulent symptoms on the *B. carinata* D5.6.12 line when subsequently tested in controlled conditions. Experiments performed in the field and in controlled conditions failed to recover a *L. maculans* population adapted to *B. carinata*.

### No evidence of gene expression of *Leptosphaeria maculans* was detected from leaves of *Brassica carinata* in field conditions

Field experiments showed that *L. biglobosa* was able to infect *B. carinata* (Fig. S1; Table 1). In order to check whether infection of *B. carinata* by *L. maculans* could be asymptomatic, we set up a transcriptomic analysis from leaves of *B. carinata* and *B. napus* cv. Darmor harvested in the field in Autumn (Fig. 1). Leaves were collected at two time points after sowing (one and two and a half months) when symptoms of *L. maculans* on plantlets of *B. napus,* sown alongside *B. carinata*, were observed. In *B. napus* leaf samples with typical *L. maculans* leaf spots (Fig. 1C), *Leptosphaeria* reads represented around 3.4% (first sampling time) to 7% (second sampling time) of the RNA-seq read numbers (Fig. 1A). In these samples, more than 97.5% of the reads were assigned to the *L. maculans* species (Fig. 1B). Although a very limited number of fungal reads were uncovered from “asymptomatic” and “atypical” *B. napus* leaves, *L. maculans* (and to a lesser extent, *L. biglobosa*) RNA-seq reads were detected, indicating that the two species were infecting these plant tissues. Even from asymptomatic samples of *B. napus*, *L. maculans* accounted for the majority of the transcriptomic activity detected, as 80-96% of the reads corresponded to *L. maculans*. In *B. napus* samples with “atypical” symptoms (Fig. 1C), both *L. maculans* and *L. biglobosa* were detected in variable proportions, with *L. maculans* reads representing 15% to 73% of the *Leptosphaeria* reads. In *B. carinata* leaf samples, a very limited amount of RNA-seq reads were assigned to either *Leptosphaeria* species (less than 1-2%; Fig. 1A) and these mostly (more than 99%) corresponded to *L. biglobosa* (Fig. 1B). These data suggest that only *L. biglobosa* can colonize *B. carinata*, even under field conditions conducive to *L. maculans* infection.

**Figure 1.**
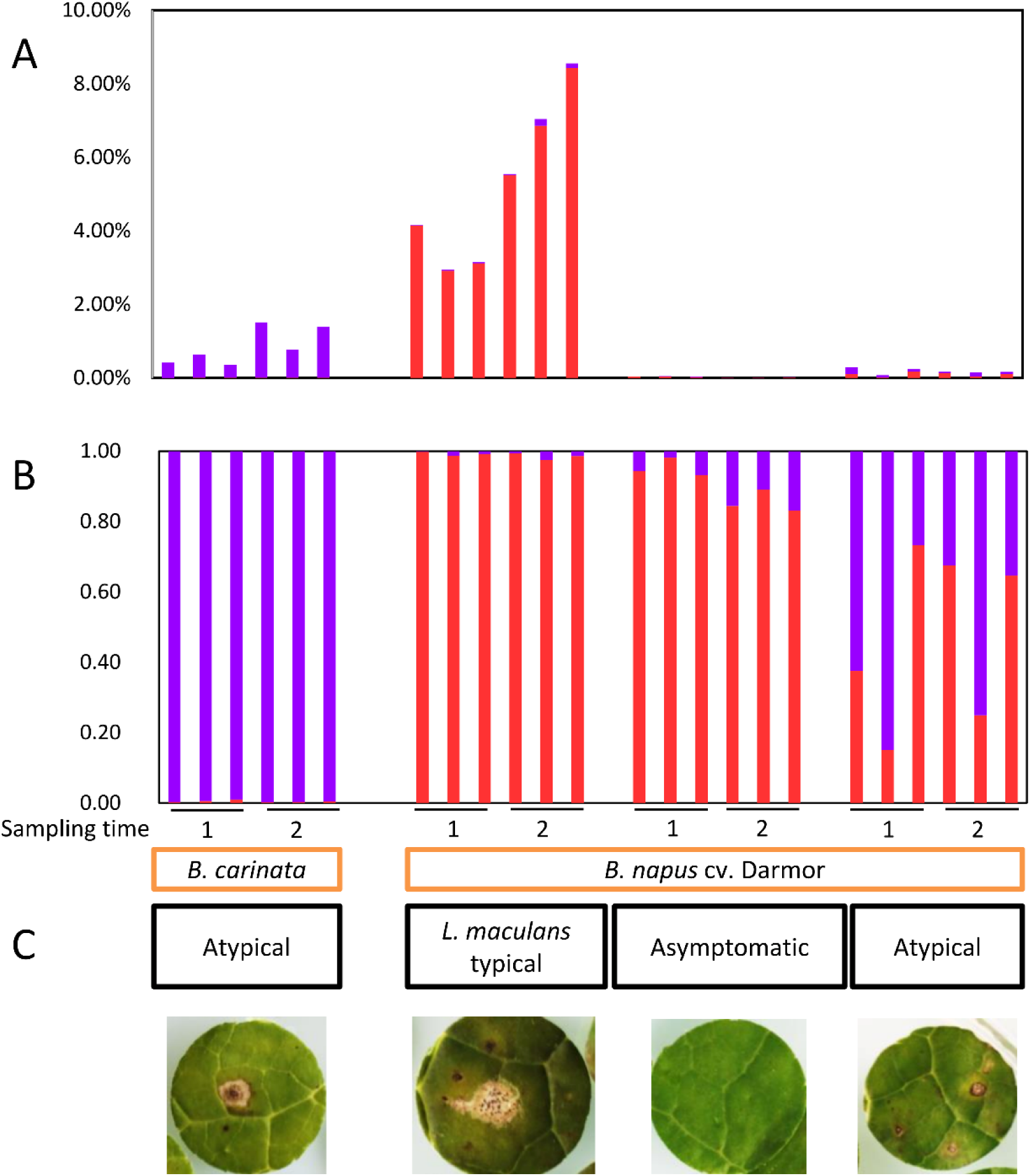
Proportion of RNA-seq reads assigned to *Leptosphaeria maculans* or *Leptosphaeria biglobosa* within field-infected leaves of *Brassica napus* (cv. Darmor) or *Brassica carinata* (line D5.6.12). (A) proportions of *L. biglobosa* (violet) and *L. maculans* (red) reads in each plant sample. (B) Among fungal reads, proportions of reads mapping to *L. biglobosa* (violet) or *L. maculans* (red) genomes. (C) Examples of symptoms observed for each type of sample sequenced. Leaf disks were collected at two sampling dates (1, two months; 2, 2.5 months after sowing); three independent replicates (different field plots) were sequenced for each time-point and plant material.

### The finding of a natural isolate of *Leptosphaeria maculans* showing partial adaptation to *Brassica carinata*

A collection of 53 isolates originating from diverse plant species and geographical origins (Table S2) were inoculated on *B. napus* cv. Westar and *B. carinata* D5.6.12 line in controlled conditions. Among them, 37 isolates originating from *B. napus* or *Raphanus sp*., and the reference isolates, JN2 and JN3, were virulent on *B. napus* cv. Westar (mean score = 4.65 ± 0.56) but failed to infect *B. carinata* (mean score = 1.05 ± 0.14), with the rapid appearance of localized necrosis directly at the point of infection (Table S2; Fig. 2A for an example of the symptom on *B. carinata* line D5.6.12). In contrast, six isolates among the 10 collected on *B. nigra* cv. Junius produced small symptoms on *B. carinata* line D5.6.12 corresponding to intermediate resistance phenotypes (mean scores ranging from 1 to 3; Table S2). These six isolates were unable to infect *B. napus* cv. Westar (mean score = 1). The isolate HB10.19, displaying an intermediate phenotype on *B. carinata* (mean score = 2.89 ± 0.6; Fig. 2A), was selected for further experimentations.

**Figure 2.**
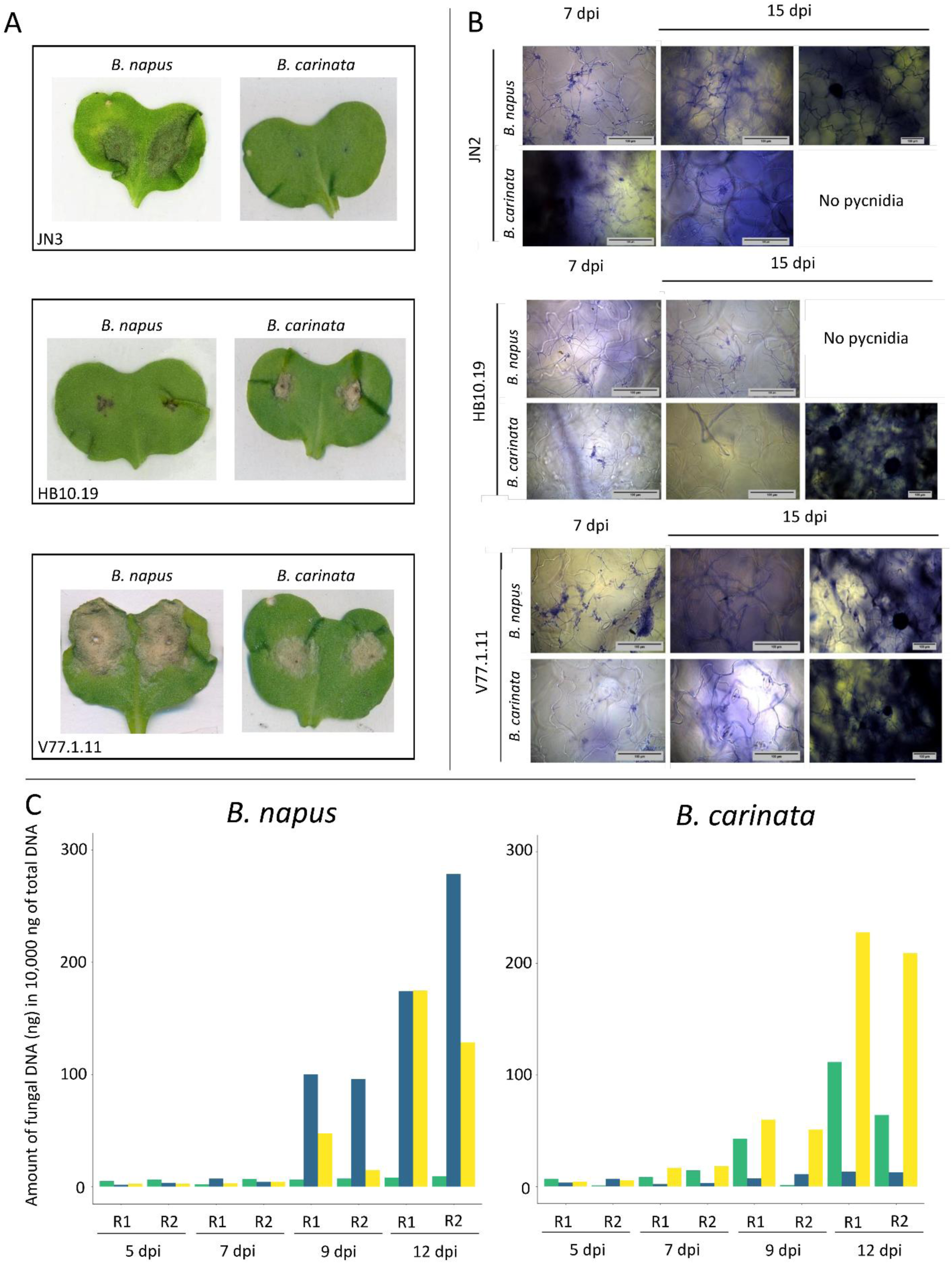
Comparison of the infection process of three *Leptosphaeria maculans* isolates on *Brassica carinata* (line D5.6.12) and *Brassica napus* (cv. Yudal). (A) Macroscopic symptoms at 12 days post inoculation (dpi) of the three isolates inoculated on *B. napus* and *B. carinata*. (B) Microscopic observation of the infection. Isolates were coloured with trypan blue. (C) Biomass quantification of the isolates. Quantification from samples recovered 5, 7, 9, and 12 dpi. Dark blue, isolate JN3; green, isolate HB10.19; yellow, isolate V77.1.11. R1, replicate 1; R2, replicate 2.

### Genetics of the interaction show a complex determinism of the adaptation of *Leptosphaeria maculans* on *Brassica carinata* and *Brassica napus*

The isolate HB10.19 was crossed with the *L. maculans* reference isolate JN3, and a progeny of 102 isolates, hereinafter referred as to “V77.i.j”, was recovered. The progeny and parental isolates were tested for pathogenicity on three plant genotypes, i.e. *B. napus* cvs. Westar, devoid of specific resistance genes, and Yudal, carrying *RlmSTEE98* (Jiquel et al., 2021) and *B. carinata* D5.6.12 line (Fig. 3). The disease scores in the progeny did not display a bimodal distribution corresponding to parental phenotypes, as expected for a monogenic inheritance of the phenotype (e.g. Balesdent *et al*., 2001), whatever the plant genotype. Instead, a continuous distribution of the mean disease scores, ranging from 1 ± 0 to 5.5 ± 0.57, was observed on *B. carinata* (Fig. 3C). A continuous distribution of the mean disease scores was also observed on the two *B. napus* cultivars tested here (Fig. 3A, B). These results suggest a polygenic determinism of the adaptation of *L. maculans* to *B. carinata* and *B. napus*.

**Figure 3.**
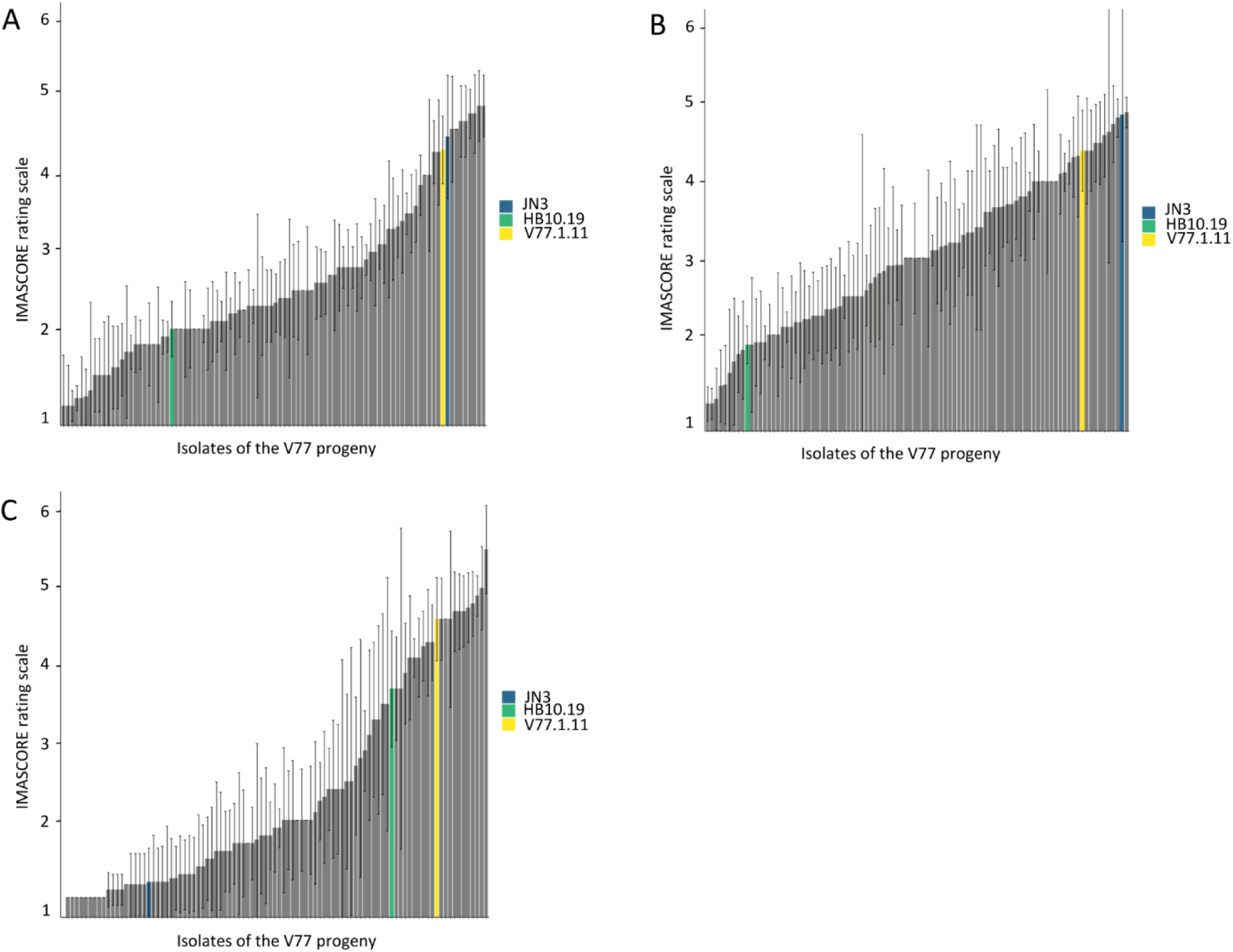
Symptoms of the V77.i.j progeny isolates showing a continuum of aggressiveness on (A) cv. Yudal, (B) cv. Westar of *Brassica napus*, and (C) *Brassica carinata* line D5.6.12. Infections and symptoms were evaluated on cotyledons following the IMASCORE rating scale (Balesdent *et al*., 2001). Dark blue bar, isolate JN3; green bar, isolate HB10.19; grey bars: V77.i.j progeny isolates, with the V77.1.11 isolate highlighted in yellow.

Among the 102 progeny isolates, some displayed a positive transgressive phenotype; i.e. they produced more severe symptoms on *B. carinata* than the parental isolate HB10.19. In addition to transgressive isolates observed on *B. carinata*, we also observed transgressive isolates on *B. napus* cv. Yudal. Four progeny isolates were identified as able to cause severe symptoms on both *B. napus* (cv. Yudal) and *B. carinata* (line D5.6.12) with symptom scores ranging from 4.6 to 5.5 on *B. carinata* and from 4 to 4.9 on *B. napus* (Fig. 3A, C). Among them, strain V77.1.11 was selected for further analysis; this progeny isolate was able to generate typical symptoms of *L. maculans* infection on cotyledons of both *B. carinata* and *B. napus*, with a greyish lesion, no black margin on *B. carinata* contrary to the parental isolate HB10.19, and extensive production of pycnidia on both species (Fig. 2A, B).

### Screening of a collection of *Brassica carinata* accessions confirmed the nonhost status of this species toward *Leptosphaeria maculans*

A collection of 52 *B. carinata* accessions recovered from the IPK genebank was screened with six isolates of *L. maculans*, including JN3, HB10.19, and the transgressive strain V77.1.11 (Table S1). All the *L. maculans* isolates failed to infect any *B. carinata* accession except for the genotype BRA_2124, which is suspected to be a cabbage based on the aspect of the cotyledons, on its susceptibility to the *L. maculans* isolates adapted to *B. napus* and on its resistance to both the parental isolate moderately adapted to *B. carinata* as well as to the transgressive isolate. Even though HB10.19 was able to moderately infect the *B. carinata* D5.6.12 line, all the IPK accessions tested here were fully resistant (with a mean score of 1.45) to this natural isolate. The transgressive progeny strain V77.1.11, induced severe susceptibility symptoms (i.e. mean score > 3) on 25 out of the 52 genotypes of *B. carinata* tested. However, a few *B. carinata* lines (n=6, mean score < 2) were resistant to V77.1.11, as infection induced a rapid local cell death typical of a HR. Conversely, some accessions were fully resistant to HB10.19 (mean score < 1.5), while they were fully susceptible (mean score > 4) to V77.1.11. As all accessions of *B. carinata* tested here were resistant to the five natural *L. maculans* isolates, this validates the extremely resistant status of *B. carinata*.

### Cotyledon-inoculations using two different approaches highlighted that only the selected progeny strain V77.1.11 gave rise to large susceptibility symptoms on *Brassica carinata*

Two inoculation approaches were performed under controlled conditions: inoculation after wounding, and inoculation with a brush, using a pycnidiospore suspension (Experimental Procedures). For inoculations after wounding, the virulence symptoms caused by JN3 on *B. napus* cv. Yudal were characterized by big grey leaf spots on which pycnidia (i.e. black dots) developed 9-14 dpi (Fig. 2A). On *B. carinata* line D5.6.12, the infection with JN3 resulted in the rapid appearance of local cell death, directly at the site of infection, typical of a HR caused by *L. maculans* on *B. napus* (Fig. 2A). The isolate HB10.19 could not infect cotyledons of *B. napus*, as displayed by the occurrence of a black necrosis, which was nevertheless more extensive than that caused by JN3 on *B. carinata* (Fig. 2A). In contrast, HB10.19 was able to infect moderately *B. carinata*, causing a grey leaf spot with pycnidia surrounded by a black margin, suggesting that the fungus can initially spread within cotyledons of *B. carinata* but is eventually blocked in its colonization. The transgressive progeny strain, V77.1.11, was capable of infecting both *Brassica* species, causing extensive grey leaf spots with pycnidia on each species (Fig. 2A). Symptoms observed following brush-inoculated cotyledons were consistent with results obtained from inoculations following wounding: on cotyledons of *B. carinata* line D5.6.12, most infections with the *L. maculans* isolate adapted to *B. napus* were followed by the appearance of local cell death (for 10 out of 16 inoculated cotyledons; Fig. S2); the isolate HB10.19 exhibited a moderate adaptation (out of 12 inoculated cotyledons, six exhibited moderate symptoms, including three showing the development of pycnidia, and six infections resulted in the development of a black symptom; Fig. S3); and most infections with V77.1.11 exhibited typically adapted phenotype (out of 18 interactions, 15 developed typical virulent symptoms, including the development of pycnidia on 12 interactions, and one black symptom was identified; Fig. S4). For brush-inoculation of *B. napus* cv. Yudal, the results were also consistent with the wounded-inoculation method, although fewer symptoms were overall observed (Fig. S5-7).

To further assess the interaction between the different isolates of *L. maculans* and *B. napus* or *B. carinata*, infection was monitored via microscopy analyses to follow the spread of the isolates in cotyledons of *B. carinata* or *B. napus*. Cotyledons of *B. napus* and *B. carinata* infected with the isolates JN2, HB10.19, or V77.1.11 were collected 7- and 15 dpi and stained with Trypan Blue (Experimental Procedures). For all interactions, pycnidiospores deposited on the surface of the cotyledon at the inoculation site after wounding were capable of germinating, and hyphae subsequently developed and were visible at 7 dpi (Fig. 2B). At 15 dpi, adapted interactions (i.e. HB10.19 infecting *B. carinata*, JN2 infecting *B. napus*, V77.1.11 infecting *B. napus* or *B. carinata*) were characterized by extensive colonization of the cotyledon, development of pycnidia (visualized by the large dark round structure) and contacts between hyphae and stmates were observed, suggesting a possible penetration of stomates, as is the case in adapted/compatible interactions. In the case of a non-adapted interaction (i.e. HB10.19 infecting *B. napus*, and JN2 infecting *B. carinata*), although limited hyphal growth was observed in the cotyledons, no pycnidia developed (Fig. 2B). The transgressive strain V77.1.11 colonized extensively both *B. napus* and *B. carinata* cotyledons, produced large amounts of pycnidia 15 dpi, and was able to penetrate stomates (Fig. 2B). Altogether, our observations showed that all isolates, adapted or non-adapted, were able to initiate the first stages of infection (i.e. pycnidiospore germination, extensive hyphal growth on the cotyledon surface) yet the non-adapted isolates failed to initiate their asexual reproduction, as is typically the case in adapted interactions, and seemed to be impaired in stomatal penetration. Although microscopic analyses show that the HB10.19 isolate was able to spread and infect successfully cotyledons of *B. carinata*, symptoms caused by this isolate on *B. carinata* were less prominent than those caused by the infection of *B. napus* by an adapted isolate, as a black margin surrounded the symptoms generated by HB10.19 (Fig. 2A).

### The natural isolate moderately adapted to *Brassica carinata,* and the progeny isolate inducing severe susceptibility symptoms on cotyledons of both *Brassica* species show a partial adaptation to *Brassica carinata*

We performed a biomass analysis to monitor fungal growth by harvesting cotyledons of *B. napus* and *B. carinata* infected by isolates JN3, HB10.19, and V77.1.11 at 5, 7, 9 and 12 dpi. On *B. napus*, the biomass of JN3 was very low 5- and 7-dpi then increased between 7- and 12-dpi to reach more than 270 ng of DNA in 10,000 ng of total DNA (Fig. 2C). Following infection of *B. carinata* by JN3, the amount of fungal material remained very low throughout the course of the experiment (the fungal biomass reached 13 ng; Fig. 2C). Likewise, the biomass of HB10.19 in infected cotyledons of *B. napus* remained low between 5- and 12-dpi (ranging between 5.35 and 9.35 ng DNA for a total DNA amount of 10,000 ng; Fig. 2C). The biomass of V77.1.11 on *B. napus* was lower than that of JN3 on the same host, although the pattern of colonization was similar to that of JN3 (a very limited biomass at 5- and 7-dpi and an increase between 7- and 12-dpi), suggesting that V77.1.11 was not able to develop as extensively as JN3 on *B. napus* (Fig. 2C). On *B. carinata*, the amount of V77.1.11 increased at 12 dpi and reached 227 ng per 10,000 ng of total DNA at the highest, suggesting this fungus is better adapted to this species than the parent HB10.19. As for the development of this isolate on *B. napus*, although the fungal biomass increased, it did not reach the same level as in the interaction between JN3 and its host. To summarize, there was an increase in fungal biomass during the adapted interactions, even though the increase was limited for interactions between HB10.19 and *B. carinata*.

We also compared stem infections between adapted and non-adapted interactions. Following inoculation of stems of *B. carinata*, no external or internal necrosis was observed with isolates JN3 and HB10.19 (Fig. 4A). For the stem infection on *B. napus*, JN3 induced external necrosis on the stem (median symptom length, 13.5 mm; Fig. 4B), and internal necrosis was systematically observed (data not shown). On the contrary, HB10.19 caused no symptoms on the outer part of the stem of *B. napus* or *B. carinata* (Fig. 4A, B). V77.1.11 caused typical adapted macroscopic symptoms on cotyledons of *B. carinata* and *B. napus* (Fig. 2A), yet limited external necrosis was observed on both *Brassica* species (Fig. 4), and inner symptoms were observed on only one of the 16 inoculated plants (data not shown). Altogether, biomass analysis and stem inoculations highlighted a limited ability to colonize *B. carinata* by HB10.19 and the transgressive strain V77.1.11.

**Figure 4.**
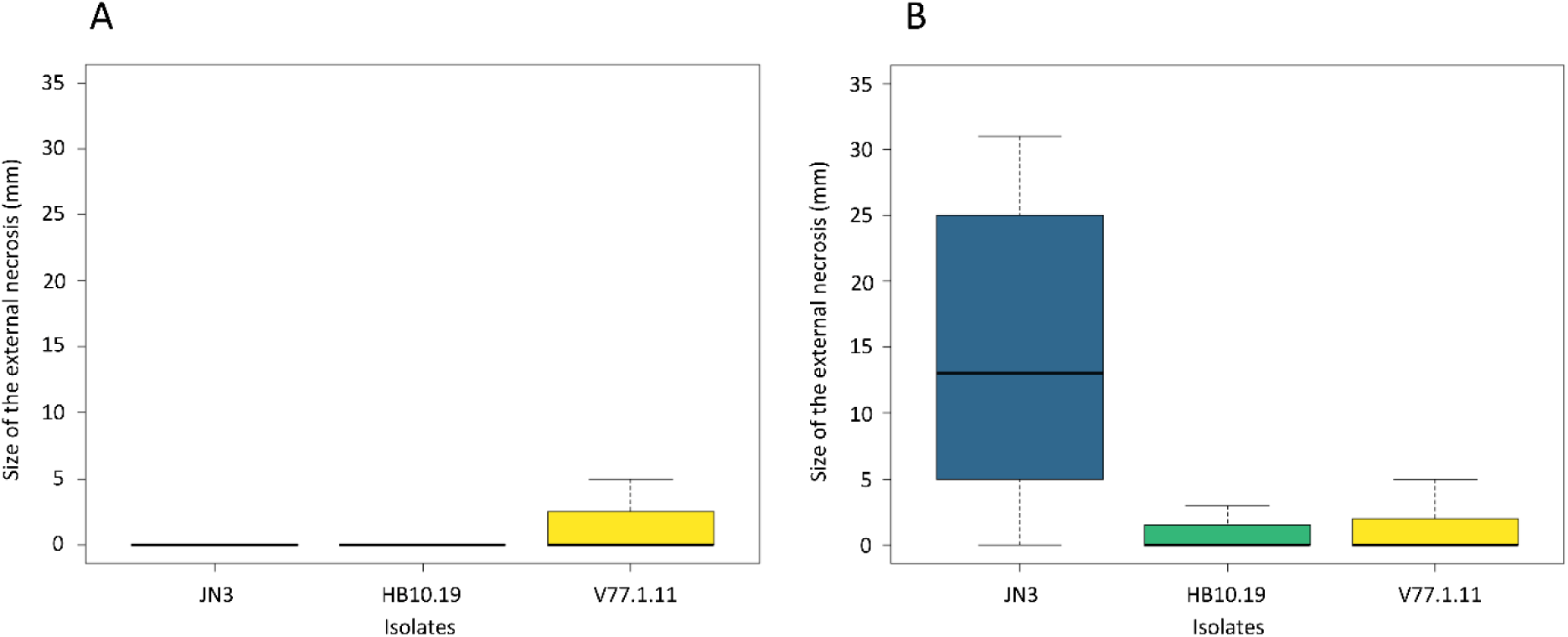
Symptoms of *Leptosphaeria maculans* on the stem basis of (A) *Brassica carinata* (line D5.6.12) and (B) *Brassica napus* (cv. Yudal).

Symptoms were evaluated by the size of the external necrosis measured in millimetre. Dark blue, isolate JN3; green, isolate HB10.19; yellow, isolate V77.1.11.

## Discussion

In this study, we highlight the extreme resistance of *B. carinata* to *L. maculans*. This result is supported by different experiments performed in the field for several years and in controlled conditions. The field experiments were performed with *B. carinata* associated with diverse *B. napus* genotypes to maximize the likelihood of contact between natural *L. maculans* isolates and *B. carinata*. Macroscopic assessment of lesions developing on *B. carinata* cotyledons and transcriptomic analyses from leaves failed to identify evidence of colonization, even asymptomatic, of *B. carinata* by *L. maculans*. We challenged the resistance of *B. carinata* in controlled conditions with different approaches (i.e. different technics of inoculation, using wounding or a brush, and different inoculum, either pycnidiospores or ascospores). We failed to recover *L. maculans* from the cotyledons of *B. carinata* with the “ascospore shower” method, although this method ensures direct ascospore ejection on the cotyledons. Moreover, although the moderately adapted isolate (HB10.19) and the transgressive adapted strain (V77.1.11) infect cotyledons of *B. carinata*, they fail to successfully infect the stem basis of *B. carinata,* showing that their adaptation to this species is only partial and restricted to specific organs. Altogether, our analyses indicate that, in our conditions, all genotypes of *B. carinata* are resistant to all the natural genotypes of *L. maculans*. Thus, we can conclude that *B. carinata* is a nonhost species of *L. maculans*.

The infection assays within the *L. maculans*/*B. napus* pathosystem in controlled conditions are routinely performed using miniaturized assays in which cotyledons are inoculated with a droplet of pycnidiospore suspension after wounding (Badawy *et al*., 1991; Mengistu *et al*., 1991). This miniaturized test is rapid and easy to handle. Nevertheless, it bypasses the natural penetration of the host via stomates. Therefore, we also performed brush inoculation to compare symptoms obtained following different inoculation methods. This assay has demonstrated that, even without wounding, *L. maculans* pycnidiospores can induce visible HR symptoms on *B. carinata*, indicating that *B. carinata* displays a type II-NHR toward *L. maculans*. In addition, during the interaction between any of the natural isolates of *L. maculans* and *B. carinata* (line D5.6.12) tested in field experiments, the rapid development of a local black necrotic symptom was identified, as observed during a hypersensitive reaction within incompatible interactions between *L. maculans* and resistant *B. napus* genotypes (Hammond and Lewis, 1987; Šašek *et al*., 2012). This response suggests that the fungus can penetrate the plant stomates but is actively recognized by the plant defence mechanism, preventing the fungus from spreading further. Type II-NHR is a post-invasive resistance characterized by a hypersensitive reaction. It is suggested to be triggered by the response to pathogen elicitors, as is effector-triggered immunity (ETI) (Gill *et al*., 2015). Type II-NHR involves active recognition and defence of the plant and relies on NB-LRR (primarily R genes) or genes involved in hormone defence pathways (Fonseca and Mysore, 2019). We postulate that *B. carinata* type II-NHR, observed during infection of natural *L. maculans* isolates could be based on the recognition of *L. maculans* effector proteins by *B. carinata* resistance proteins. Moreover, the effector repertoire is one of the key determinants of pathogen host range, including adaptation to different host genotypes or host species (Ma *et al*., 2010; Raffaele *et al*., 2010; Spanu *et al*., 2010; Dong *et al*., 2015; Frantzeskakis *et al*., 2018). In *L. maculans*, the annual sexual reproduction favours rising of avirulence loci via recombination (Rouxel and Balesdent, 2017). The continuum of aggressiveness caused by the V77 progeny strains on *B. carinata* suggests that the rearrangement of traits, including effectors, between isolates JN3 and HB10.19 allows the progeny strains to be better adapted to *B. carinata*. These progeny strains may lack effectors allowing them to escape the plant recognition. The advantage of type II-NHR is that plants displaying such resistance can be used to identify control methods against pathogens. Among the *B. carinata* genotypes recovered from the IPK and tested here, we identified a few displaying susceptibility towards the transgressive strain V77.1.11 while all are resistant to the parental isolates used for the cross. Understanding the mechanisms underlying the extreme resistance of *B. carinata* will enable identification of new sources of resistance that may be more durable than the host resistances against stem canker usually used in breeding programs.

On the contrary, Type-I NHR prevents fungus development, as the pathogen can be stopped directly upon contact between the non-adapted pathogen and the nonhost species by the inability of the inoculum to germinate, absence of haustoria development following germination or inability to penetrate (i.e. Cheng *et al*., 2012, 2013; Plotnikova *et al*., 2023). In the *B. carinata*/*L. maculans* interaction, microscopic analysis evidences that pycnidiospores germinate at the surface of *B. carinata* cotyledons both in the adapted and the non-adapted interactions. In adapted interactions, contact between hyphae and stomates were observed, suggesting penetration of *B. carinata* stomates, as is the case for adapted isolates on *B. napus*. Yang and Fernando (2021) performed microscopic analyses between various isolates of *L. maculans* and different genotypes of *B. napus,* showing that an adapted interaction is associated with hyphal growth, colonization of intercellular spaces, and pycnidia formation, consistent with previous analyses (Hammond *et al*., 1985; Li *et al*., 2008). In contrast, neither pycnidia formation nor extensive necrosis was observed in the non-adapted interactions, as in an incompatible interaction. Chen and Howlett (1996) showed that for an incompatible interaction between *L. maculans* and *B. napus*, guard cell necrosis and tissue browning appear when hyphae have reached the mesophyll layer. We observe here that *L. maculans* is still viable on the surface of cotyledons in non-adapted interactions, however, further microscopic analyses are required to better characterize the nonhost interaction between *L. maculans* and *B. carinata* and to identify when the infection process is impaired.

Among the 53 natural isolates tested on *B. carinata* in controlled conditions, we identified four isolates from our collection that could moderately infect one line; among these, HB10.19 was selected for further analysis. This isolate was recovered from *B. nigra* leaves in the field. *B. carinata* results from natural hybridization between *B. oleracea* and *B. nigra*, ca. 11,000-29,900 years ago (Yim *et al*., 2022); hence *B. nigra* and *B. carinata* share the B-genome. It has been demonstrated that the B-genome is an important resource for high level of resistances against biotic and abiotic stresses, including stem canker (Christianson *et al.,* 2006; Fredua-Agyeman *et al.,* 2014; Rashid *et al.,* 2018; Yim *et al.,* 2022). Further analysis of the *B. carinata* genome will undoubtedly pinpoint genomic features associated with this resistance.

Interestingly, all isolates recovered on *B. carinata* leaves from the field experiment corresponded to *L. biglobosa*. *L. maculans* and *L. biglobosa* are adapted to Brassicaceae species, and share common epidemiological, infection strategies and ecological niches, thus can be found co-infecting oilseed rape (West *et al*., 2001; Mendes-Pereira *et al*., 2003; Jacques *et al*., 2021). The divergence between *L. maculans* and *L. biglobosa* is estimated at 22 MYA (Grandaubert *et al*., 2014). Despite their similar life cycles, *L. biglobosa* and *L. maculans* have genomic specificities, such as the fact that *L. biglobosa* has more putative genes encoding secondary/specialized metabolites, there are different effector repertoires between the two fungal species, and a massive expansion of transposable elements in the *L. maculans* genome (Grandaubert *et al.,* 2014). These genomic characteristics could explain their different adaptive traits, as already hypothesized by Grandaubert *et al*. (2014), suggesting a link between the impact of the different genomic features of the *L. maculans*/*L. biglobosa* species complex on their adaptation.

To investigate the extreme resistance of *B. carinata* toward *L. maculans*, we performed a cross between a reference isolate JN3, adapted to *B. napus*, and the isolate HB10.19, moderately adapted to *B. carinata*. In the offspring of the cross, several transgressive strains were identified, showing greater aggressiveness on one of the two *Brassica* species than the parental isolate adapted to this *Brassica*. Extreme phenotypes within progenies have critical roles in species evolution (Rieseberg *et al*., 1999). Four progeny isolates were able to cause symptoms on both *Brassica* species. These transgressive phenotypes were also observed when we quantified the amount of biomass of the three *L. maculans* isolates JN3, HB10.19, and V77.1.11 on *B. napus* and *B. carinata*. Although V77.1.11 produces more biomass than HB10.19 on *B. carinata* during infection, this biomass does not reach the biomass of *L. maculans* adapted to *B. napus*. Transgression segregation has been well described in plants (Rieseberg *et al*., 1999, 2003; Stelkens and Seehausen, 2009), nevertheless, to date, only a few transgressive phenotypes were formally laid out in filamentous fungi (Cumagun *et al*., 2004; Voss *et al*., 2010; Gibson *et al*., 2014; Caffier *et al*., 2022). The transgressive isolates obtained in the progeny indicate that after only one sexual reproduction event, it is possible to generate strains of *L. maculans* adapted to *B. carinata* cotyledons. We hereby demonstrate that the adaptation of *L. maculans* to *B. carinata* is under a polygenic determinism. Our present study represents a valuable biological material to investigate underlying factors of the adaptation of *L. maculans to different Brassica species. Pathogens that have sexual and asexual reproduction, large population size, and high mutation rate are very likely to evolve rapidly and bypass plant host-resistance (McDonald and Linde, 2002). L. maculans is a fungus that displays both types of reproduction mode within a single lifecycle (Rouxel and Balesdent, 2005), a large population of isolates containing genetic polymorphism (Rouxel et al., 2003; Balesdent et al., 2006, 2022) and a high recombination rate, mainly localized in GC-rich isochores (Rouxel et al., 2011) which makes this fungus a rapidly evolving species and a model of interest to study adaptation and evolution. The transgressive strains and the cross between JN3 and HB10.19 will allow us to identify the genetic mechanisms underlying adaptation of L. maculans to different hosts and identify whether adaptation to nonhost resistance of B. carinata involves the same underlying molecular events as those involved in the host adaptation*.

Despite NHR being considered more durable than host resistance (Ayliffe and Sørensen, 2019; Fonseca and Mysore, 2019), it is not insurmountable. Even though *B. carinata* is not a widely cultivated species, it has been introduced into North America and Argentina for industry uses (Taylor *et al*., 2010; Ban *et al*., 2017; Song *et al*., 2021). The sympatry of *B. napus* and *B. carinata* may favour a host shift of *L. maculans*, or an expansion of its host range, toward *B. carinata*. Several pathogens have adapted due to the proximity of their host with another plant species (Bettgenhaeuser *et al*., 2014; Xia *et al*., 2018). Consequently, the risk of emergence of a *L. maculans* population adapted to *B. carinata* must be considered if oilseed rape crops are grown alongside *B. carinata* crops. Besides, environmental conditions (i.e., temperature, photoperiod, humidity, developmental stage of the plant, etc.) need to be considered and can affect the nonhost status of a plant toward pathogens (Panstruga and Moscou, 2020; Nnadi and Carter, 2021). The context of global climate change represents an increasing threat of the emergence of new and better-adapted pathogen species (for examples, see Nnadi and Carter, 2021). Thus, global climate changeould also create conditions favouring host shifts in *L. maculans*.

Altogether, our analyses show that the *L. maculans*/*B. carinata* pathosystem is a promising nonhost framework to decipher the underlying mechanisms responsible for the adaptation of *L. maculans* to different hosts, and to investigate the genetic determinants involved in the NHR of *B. carinata* toward *L. maculans*. Nevertheless, we highlighted the risk of the emergence of better-adapted isolates to this *Brassica* species, which might be emphasized by agricultural practices and climate change.

## Author contributions

MHB and JLS supervised the work; TR, MHB and JLS conceptualized the experiments; TR, MHB, JLS funding acquisition; JMN, MG, CAR, EJG, TR, MHB, JLS performed experiments; JMN, MG, CAR, EJG, TR, MHB, JLS: validation; JMN, CAR, EJG, MHB, JLS: visualization; JMN, EJG, TR, MHB, JLS drafted the paper, all authors revised and approved the final version of the paper.

## Acknowledgments

This work was founded by the French National Research Agency project AvirLep (ANR GPLA07-024C) and a grant from the INRAE department “Santé des plantes et environnement (SPE)” (Project Phytoadapt). Julie M. Noah benefited from a grant by INRAE (SPE) and the “Club Phoma” consortium, Mathilde Gorse benefited from a grant by the Saclay Plant Sciences-SPS (ANR-17-EUR-0007). The authors particularly thank partners of the Avirlep project for field assays and collection of infected leaves; the experimental unit at INRAE (Grignon) for the management of the fields at Grignon; Laurent Coudard (BIOGER) for isolate management; the greenhouse technician staff for plant management; the administrative supporting staff for the administrative and financial follow-up of this project, and the laboratory glassware staff of the BIOGER research unit. We thank Dr. Joao Silva-Dias (Technical University of Lisbon) for the *B. carinata* line D5.6.12, Evelin Willner, from the IPK Gatersleben, for providing us with seeds of various genotypes of *B. carinata,* INRAE-IGEPP and the BrACySol CRB (INRAE) for the Fu-Eurol-*Rlm10* genotype, Hortense Brun (IGEPP) and Dr. Martin Barbetti (University of Western Australia) for selected isolates. The “Effectors and Pathogenesis of *L. maculans*” (EPLM) group benefits from the support of Saclay Plant Sciences-SPS (ANR-17-EUR-0007).

## Supplementary tables

**Table S1. Accessions of *Brassica carinata* retrieved from the IPK genebank and tested for their resistance toward different isolates of *Leptosphaeria maculans*.**

^a^acquisition date from the IPK Genebank;

^b^country of origin; ETH, Ethiopia; ZMB, Zambia; SWE, Sweden; PAK, Pakistan;

^c^mean of the symptoms assessed using the IMASCORE rating scale (Balesdent *et al*., 2001);

^d^origin of the *L. maculans* isolates: HB10.19 and V77.1.11, this study; IBCN14 and IBCN80 are reference isolates of *L. maculans* included in the International Blackleg of Crucifer Network collection (Balesdent *et al*., 2005).

**Table S2. List and characteristics of *Leptosphaeria maculans* isolates used and their pathogenicity on *Brassica napus* and *Brassica carinata*.**

^a^Symptoms were evaluated on 10-12 plantlets of *B. napus* and *B. carinata* and were evaluated as described in Balesdent *et al*., 2001, with a score of 1-3 corresponding to an avirulent isolate and a resistant plant and scores ranging from 4 to 6 corresponding to a virulent isolate and a susceptible plant, according to the IMASCORE scale.

**Table S3. Primers used for PCR and qPCR amplification in this study.** For, forward primer, and Rev, reverse primer. Probes are used to improve the specific quantification of a species (Jacques *et al*., 2021).

**Figure S1.**
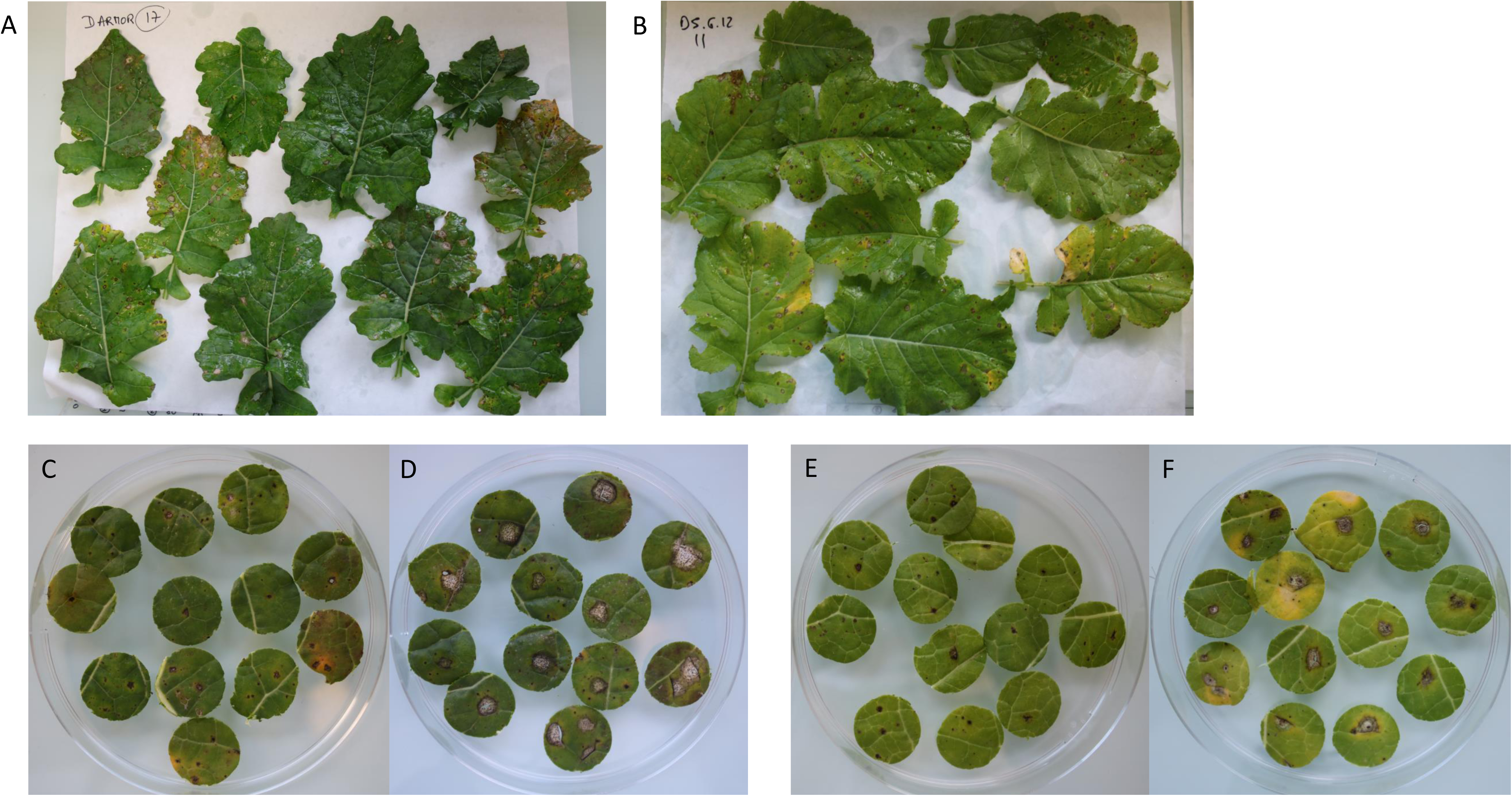
Symptoms recovered on leaves of *Brassica napus* (cv. Darmor) and *Brassica carinata* (line D5.6.12) in a field experiment. Leaves of (A) *B. napus* cv. Darmor and (B) *B. carinata* line D5.6.12. Examples of symptoms observed on leaf disks collected 2.5 months after sowing on (C), (D) *B. napus* and (E), (F) *B. carinata*. Symptoms identified from disks (C), and (E) were suspected to be caused by *L. biglobosa*; while (D) and (F) were suspected to be caused by *L. maculans*.

**Figure S2.**
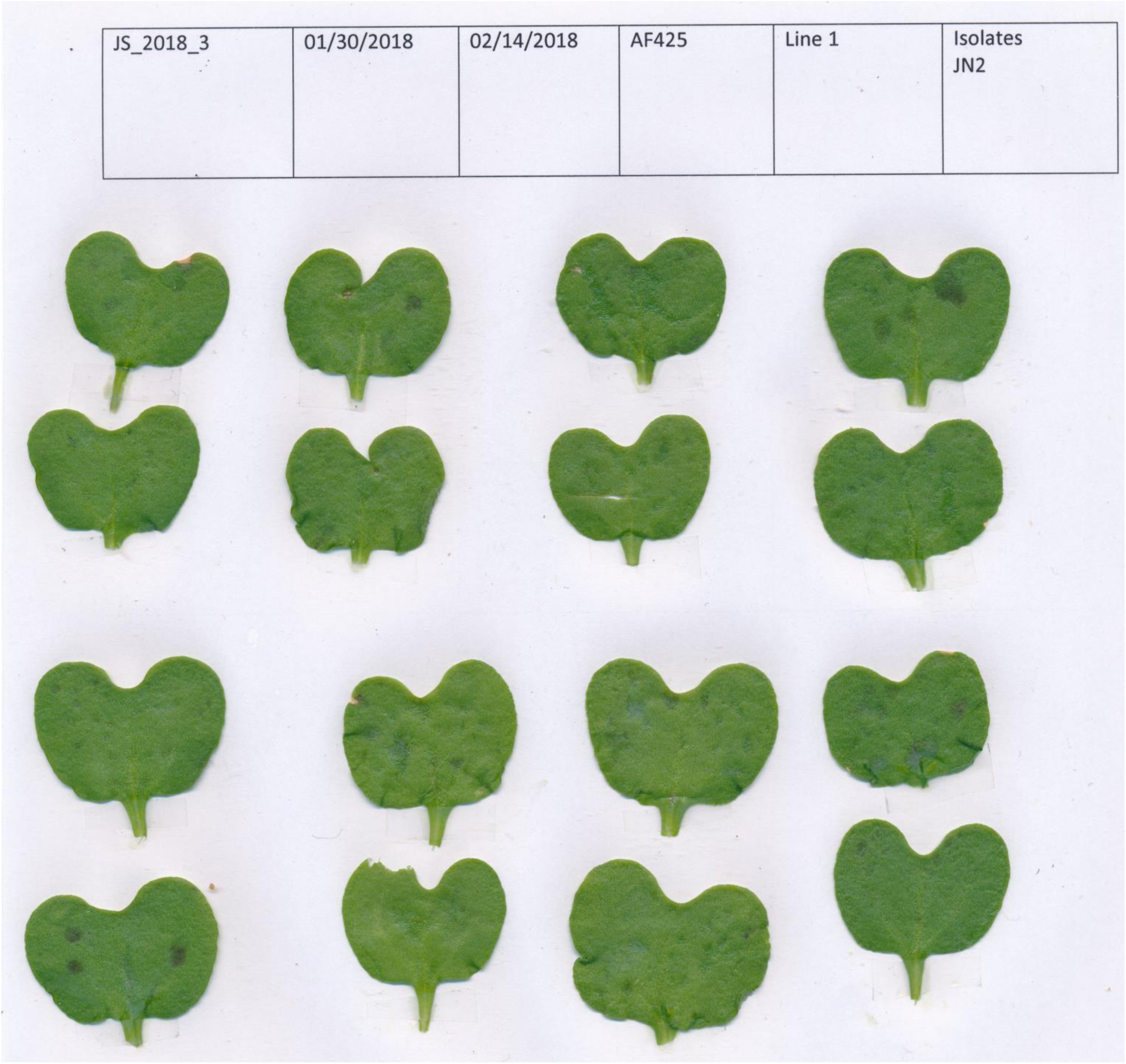
Symptoms observed on *Brassica carinata* line D5.6.12 following brush inoculation with the isolate JN2 of *Leptosphaeria maculans*. The upper line displays details of the experiment; with, from left to right, JS_2018_3, the reference of the inoculation; the dates of start and end of the experiment (i.e. dates of inoculation and observation); AF425, the reference of the *B. carinata* line used for the inoculation (i.e. line D5.6.12); Line 1, the line of the plants used for inoculation; JN2, the name of the *L. maculans* isolate brush-inoculated on the cotyledons (Experimental Procedures).

**Figure S3.**
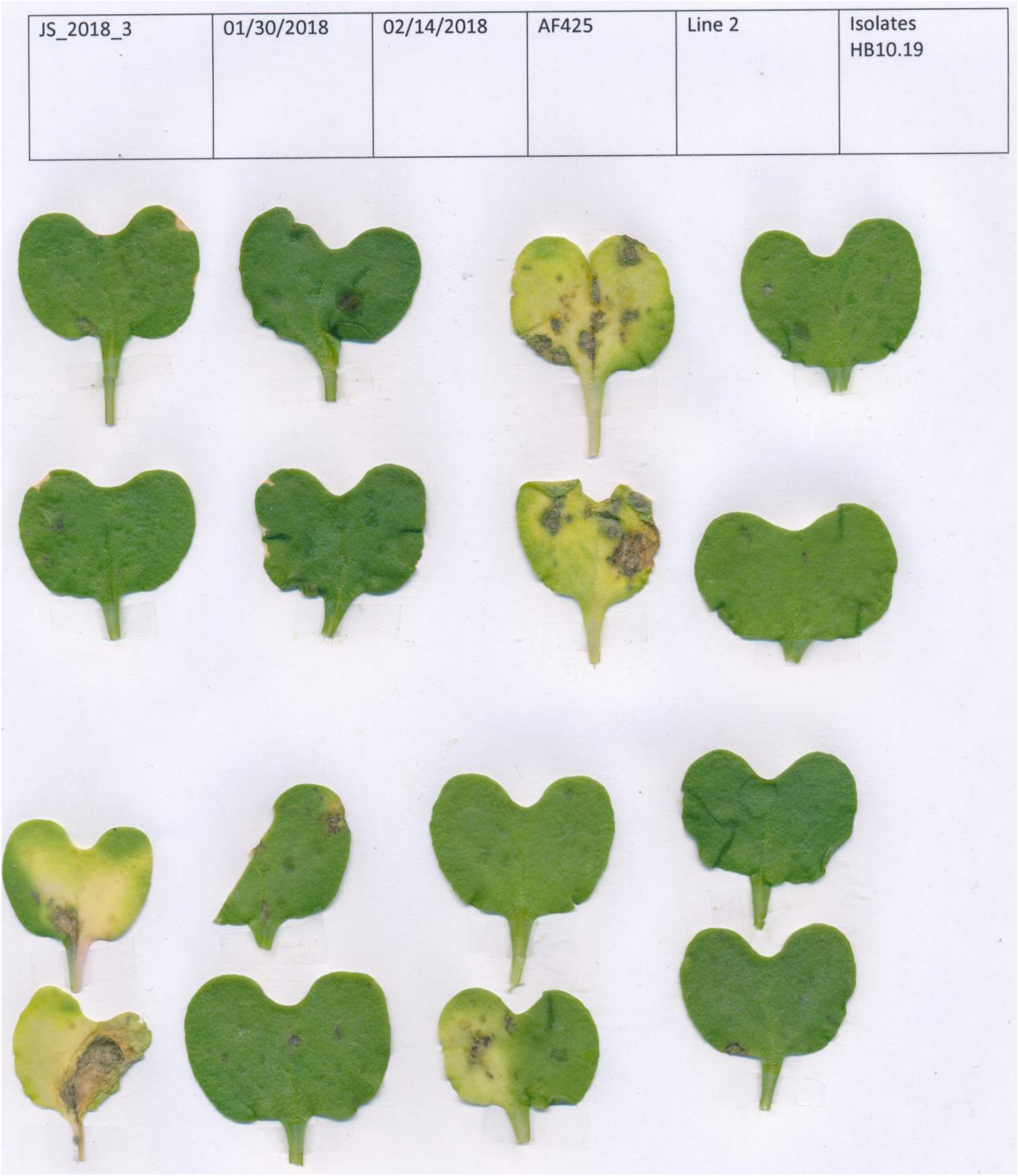
Symptoms observed on *Brassica carinata* line D5.6.12 following brush inoculation with the isolate HB10.19 of *Leptosphaeria maculans*. The upper line displays details of the experiment; with, from left to right; JS_2018_3, the reference of the inoculation; the dates of start and end of the experiment (i.e. dates of inoculation and observation); AF425, the reference of the *B. carinata* line used for the inoculation (i.e. line D5.6.12); Line 2, the line of the plants used for inoculation; HB10.19, the name of the *L. maculans* isolate brush-inoculated on the cotyledons (Experimental Procedures).

**Figure S4.**
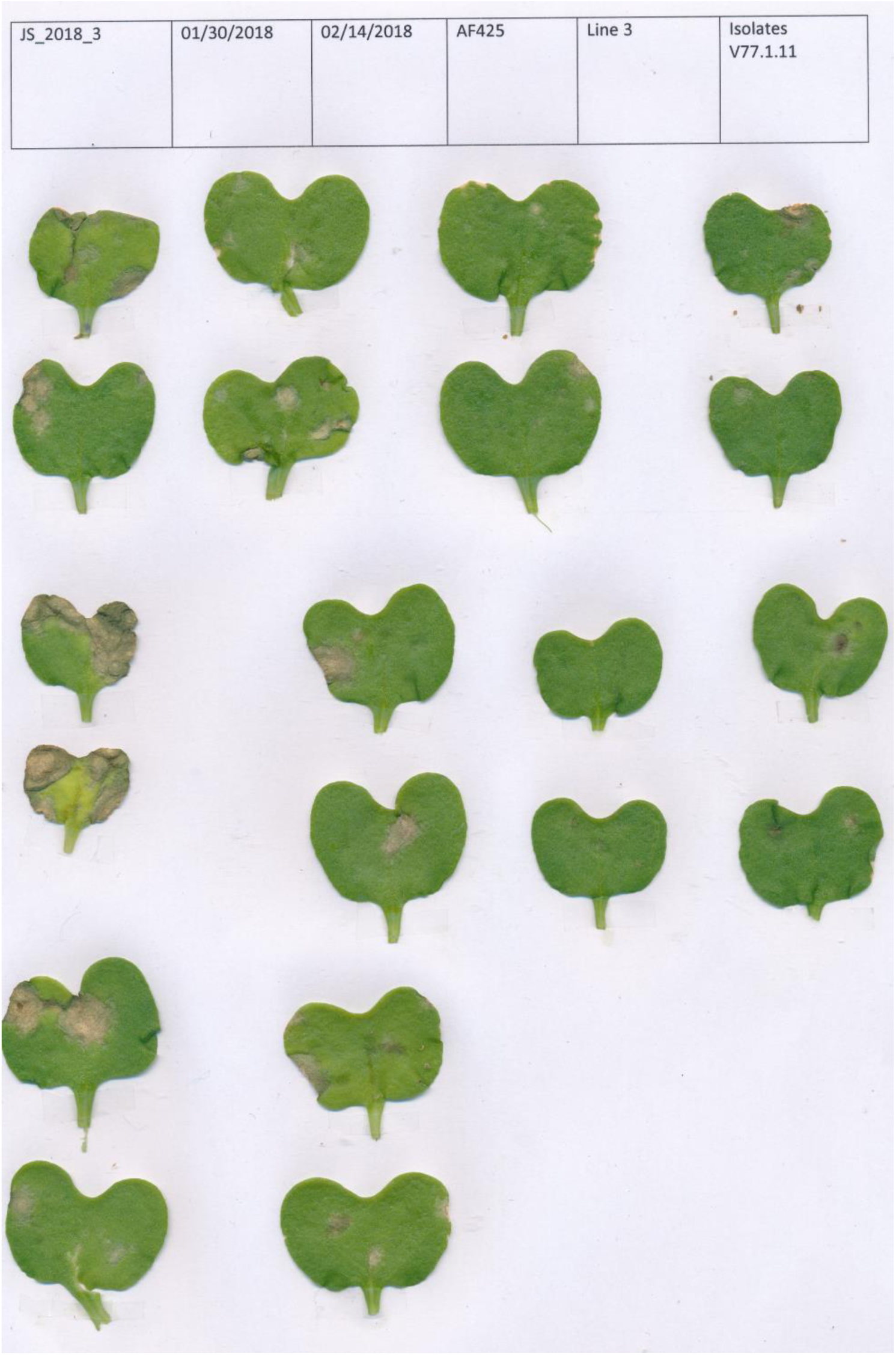
Symptoms observed on *Brassica carinata* line D5.6.12 following brush inoculation with isolate V77.1.11 of *Leptosphaeria maculans*. The upper line displays details of the experiment; with, from left to right; JS_2018_3, the reference of the inoculation; the dates of start and end of the experiment (i.e. dates of inoculation and observation); AF425, the reference of the *B. carinata* line used for the inoculation (i.e. line D5.6.12); Line 3, the line of the plants used for inoculation; V77.1.11, the name of the *L. maculans* isolate brush-inoculated on the cotyledons (Experimental Procedures).

**Figure S5.**
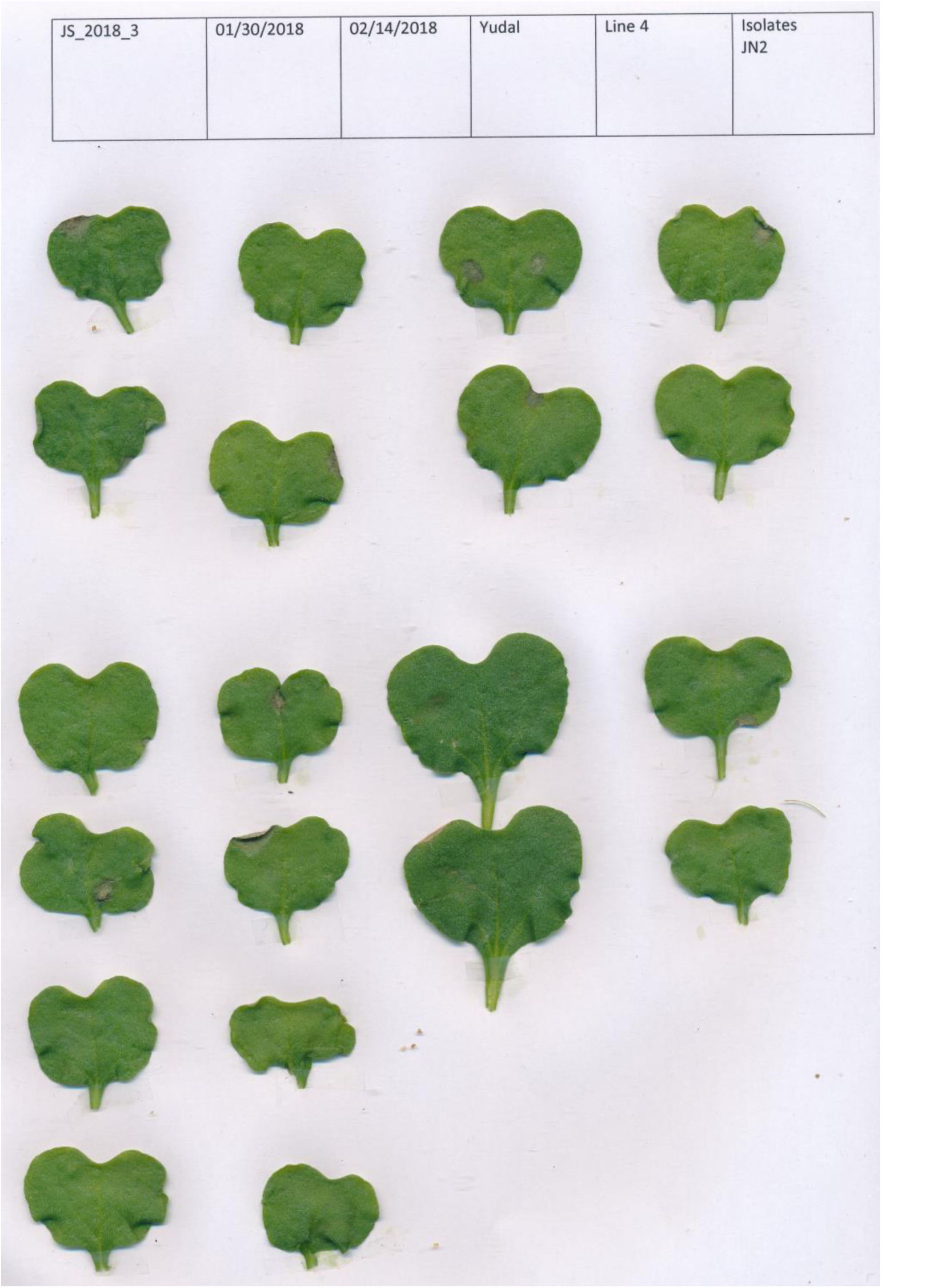
Symptoms observed on *Brassica napus* cv. Yudal following brush inoculation with the isolate JN2 of *Leptosphaeria maculans*. The upper line displays details of the experiment; with, from left to right; JS_2018_3, the reference of the inoculation; the dates of start and end of the experiment (i.e. dates of inoculation and observation); Yudal, the reference of the *B. napus* cultivar used for the inoculation; Line 4, the line of the plants used for inoculation; JN2, the name of the *L. maculans* isolate brush-inoculated on the cotyledons (Experimental Procedures).

**Figure S6.**
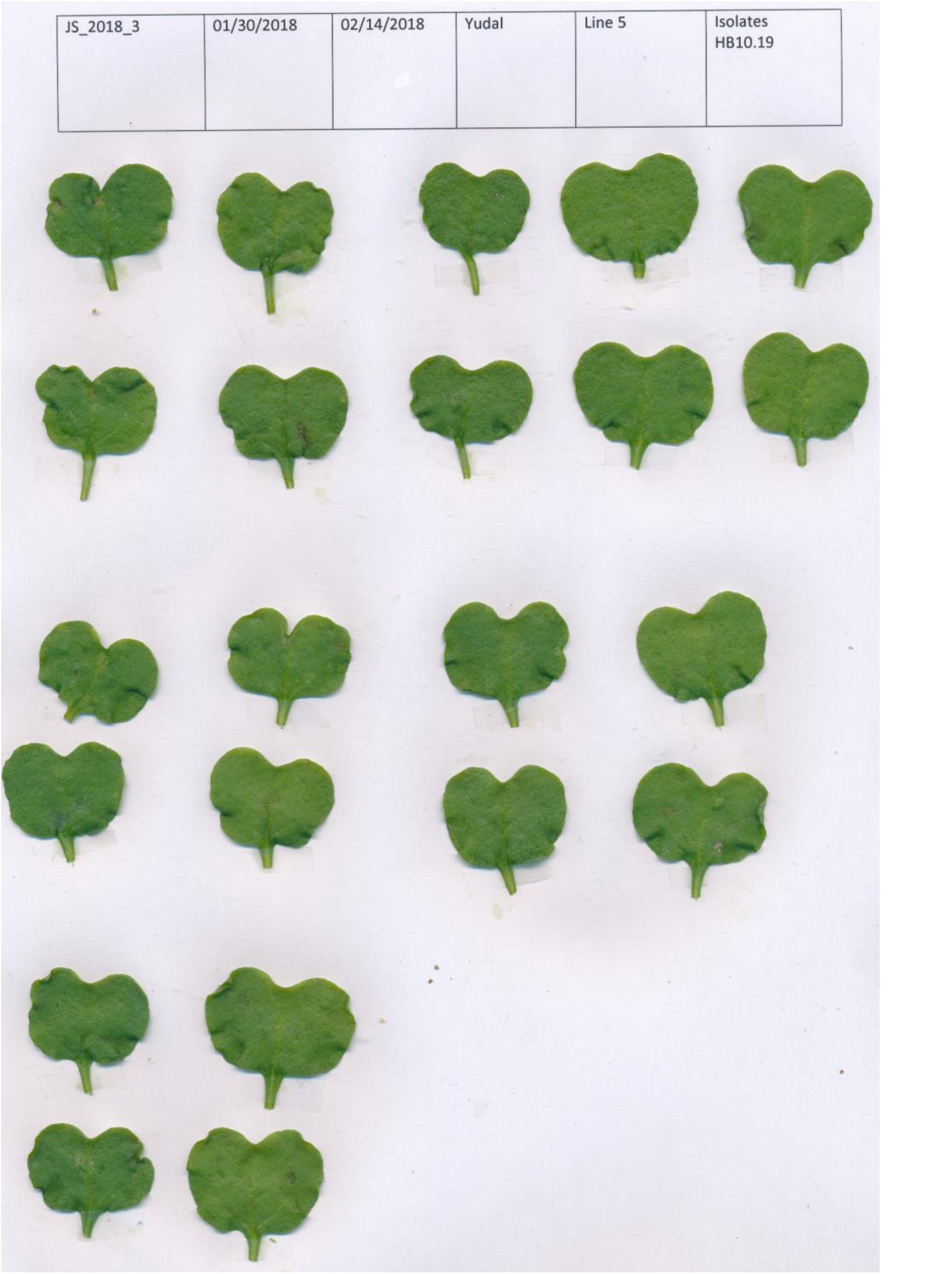
Symptoms observed on *Brassica napus* cv. Yudal following brush inoculation with the isolate HB10.19 of *Leptosphaeria maculans*. The upper line displays details of the experiment; with, from left to right; JS_2018_3, the reference of the inoculation; the dates of start and end of the experiment (i.e. dates of inoculation and observation); Yudal, the reference of the *B. napus* cultivar used for the inoculation; Line 5, the line of the plants used for inoculation; HB10.19, the name of the *L. maculans* isolate brush-inoculated on the cotyledons (Experimental Procedures).

**Figure S7.**
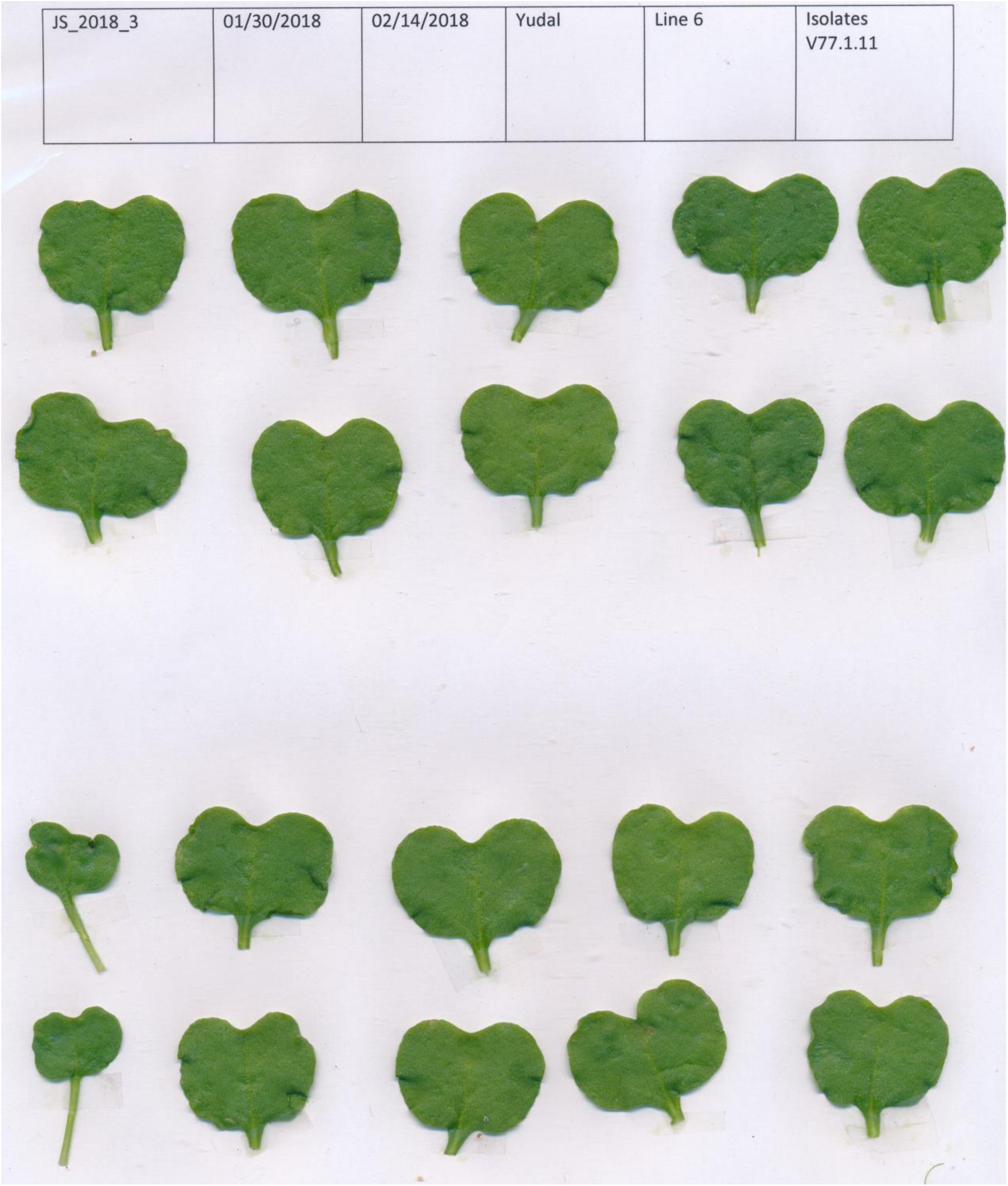
Symptoms observed on *Brassica napus* cv. Yudal following brush inoculation with the isolate V77.1.11 of *Leptosphaeria maculans*. The upper line displays details of the experiment; with, from left to right; JS_2018_3, the reference of the inoculation; the dates of start and end of the experiment (i.e. dates of inoculation and observation); Yudal, the reference of the *B. napus* cultivar used for the inoculation; Line 6, the line of the plants used for inoculation; V77.1.11, the name of the *L. maculans* isolate brush-inoculated on the cotyledons (Experimental Procedures).

